# The inner nuclear membrane protein NEMP1 is required for nuclear envelope openings and enucleation of erythroblasts during erythropoiesis

**DOI:** 10.1101/2022.05.26.493585

**Authors:** Didier Hodzic, Jun Wu, Karen Krchma, Andrea Jurisicova, Yonit Tsatskis, Yijie Liu, Peng Ji, Kyunghee Choi, Helen McNeill

## Abstract

Nuclear Envelope Membrane Proteins (NEMP) are a conserved family of nuclear envelope proteins that reside within the inner nuclear membrane. Even though *Nemp1* knockout (KO) mice are overtly normal, they display a pronounced splenomegaly. This phenotype and recent reports describing a requirement for nuclear envelope openings during erythroblasts terminal maturation led us to examine a potential role for Nemp1 in erythropoiesis. Here, we report that *Nemp1* knockout (KO) mice show peripheral blood defects, anemia in neonates, ineffective erythropoiesis, splenomegaly and stress erythropoiesis. The erythroid lineage of *Nemp1* KO mice is overrepresented until the pronounced apoptosis of polychromatophilic erythroblasts. We show that NEMP1 localizes to the nuclear envelope of erythroblasts and their progenitors. Mechanistically, we discovered that NEMP1 accumulates into aggregates that localize near or at the edge of nuclear envelope openings and *Nemp1* deficiency leads to a marked decrease of both nuclear envelope openings and ensuing enucleation. Together, our results for the first time demonstrate that NEMP1 is essential for nuclear envelope openings and erythropoietic maturation *in vivo* and provide the first mouse model of defective erythropoiesis directly linked to the loss of an inner nuclear membrane protein.

## INTRODUCTION

<u>N</u>uclear <u>E</u>nvelope <u>M</u>embrane <u>P</u>rotein1 (NEMP1, encoded by *Tmem194a*) is a highly conserved multipass transmembrane protein that resides within the inner nuclear membrane (INM) of the nuclear envelope (NE). We recently showed that genetic inactivation of *Nemp1* leads to a loss of fertility in worm, fish, and flies [1]. In mice, *Nemp1* is required for female fertility but dispensable for male fertility [1]. Except for the presence of a conserved domain of unknown function (DUF 2215) that encompasses its transmembrane and proximal nucleoplasmic C-terminal region, NEMP1 does not harbor any known functional motif. However, its nucleoplasmic region has been shown to interact with barrier to autointegration factor (BAF) and RAN GTPase to mediate *Xenopus* eye development [2,3]. NEMP orthologs have recently been identified in plants (PNET2a, b and c) where they play an essential role in chromatin architecture [4]. Using BioID as well as affinity purification followed by mass spectrometry, we recently showed that NEMP1 interacts with LEM domain proteins EMERIN, MAN1 and LAP2, known to physically link the NE to chromatin and support mechanical stiffness. Accordingly, we showed that loss of *Nemp1* expression drastically affects NE mechanical stiffness in cultured cells and oocytes [1].

Mammalian erythropoiesis consists of the differentiation of hematopoietic stem cells into megakaryocyte-erythrocyte progenitors (MEPs) that generate burst forming unit-erythroid (BFU-E) that in turn differentiate into colony forming unit-erythroid (CFU-E). The latter generate proerythroblasts (ProE) that correspond to the first recognizable erythroid cell. During terminal erythropoiesis, ProE undergo 4-5 mitoses that generate basophilic (EryA), polychromatophilic (EryB) and orthochromatic (EryC) erythroblasts. Erythroblast differentiation is characterized by chromatin condensation which is required for enucleation, the ultimate step of erythropoiesis that generates pyrenocytes and reticulocytes [5–8]. Interestingly, recent studies have established that recurrent NE openings in maturing erythroblasts allow for the partial and selective release of histones in the cytoplasm, a biological process that is essential for chromatin condensation and final enucleation[9–12]. However, the role of NE proteins in this remarkable biological process remains to be established.

Adult *Nemp1* knockout (KO) mice are overtly normal. However, both *Nemp1* KO males and females display strikingly enlarged spleens. This phenotype and the involvement of NE openings in terminal erythropoiesis led us to examine the biological function of *Nemp1* in erythropoiesis. We show that *Nemp1* KO mice display erythroid lineage differentiation defects. Polychromatophilic erythroblasts displayed reduced frequencies of NE openings and of enucleation as well as increased apoptosis, leading to erythroid maturation defects. These data show that NEMP1 is required for NE openings and enucleation during the late stages of erythroblast maturation.

## RESULTS

### *Nemp1* KO mice have splenomegaly and abnormal erythropoiesis

*Nemp1* KO mice displayed significantly enlarged spleens with increased cellularity compared to heterozygous *Nemp1* or wild type (WT) mice (Figure 1A, B, C). Wright-Giemsa staining of *Nemp1* KO blood smears showed red blood cells (RBCs) with irregular shapes and spiky membranes (Figure 1D, red arrows). Complete blood count (CBC) analyses of peripheral blood (PB) using a Hemavet further showed decreased RBC counts in neonates suggestive of anemia although adult mice showed no such obvious phenotype (Figure 1E and S1). RBCs showed decreased hemoglobin content and higher red cell distribution widths (RDW), reflecting irregular RBC membrane shapes, two phenotypes that persisted throughout life (Figure 1E and S1). Finally, FACS analysis of PB also showed increased percentage of immature RBCs in the circulation (Figure 1F). The bone marrow (BM) of *Nemp1* KO mice also appeared more densely packed and displayed increased cellularity by comparison to WT BM (Figure 1G).

**Figure 1:**
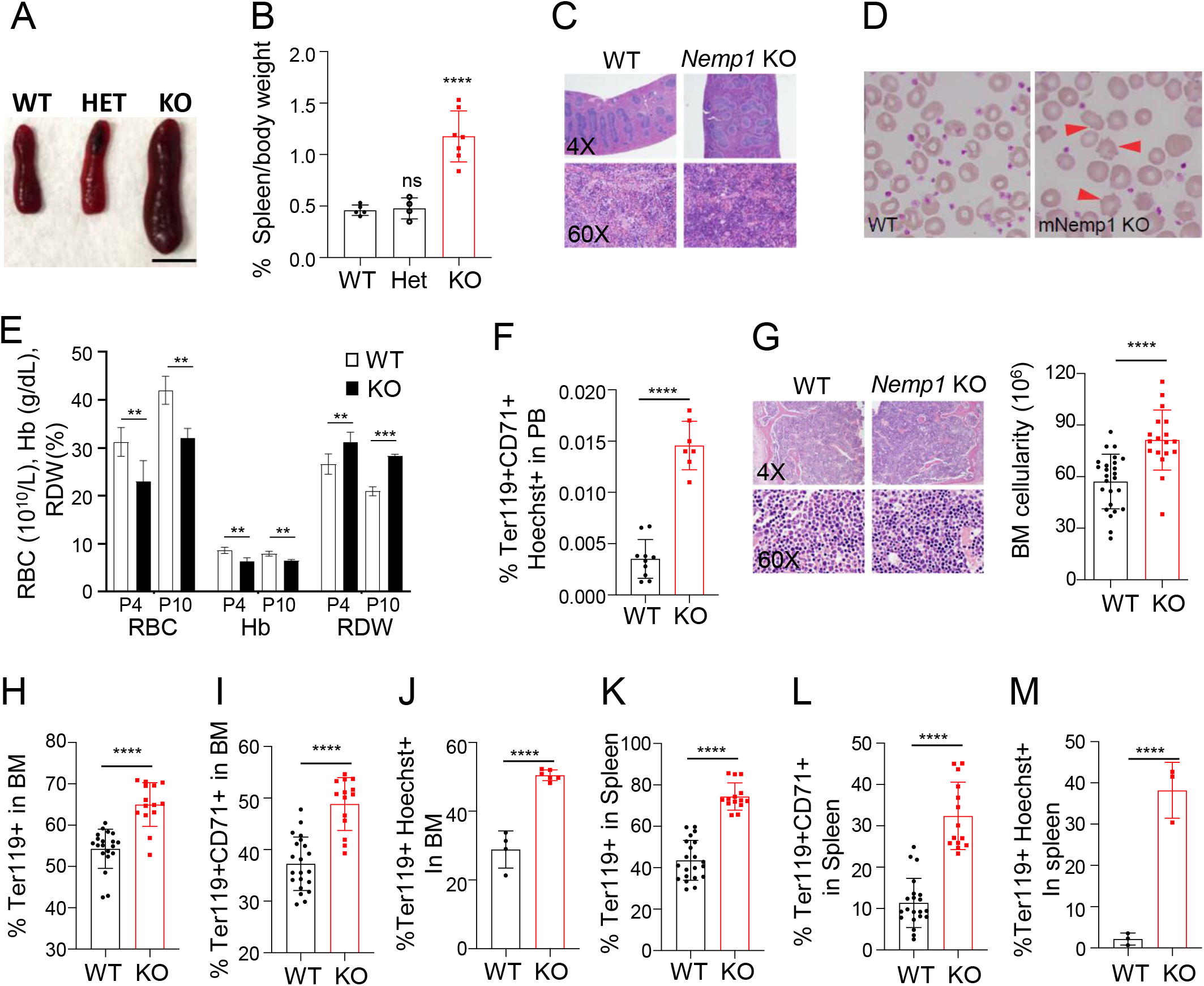
Nemp1 KO mice display splenomegaly, blood defects and erythroid lineage expansion. Representative spleen size (**A**) and spleen/body weight ratios (**B**) across genotypes. Scale bar: 1 cm. **C.** H&E staining of WT and Nemp1KO spleen. **D**. Wright-Giemsa staining of WT and Nemp1 KO blood smears. Red arrows point to cytoplasmic membrane spikes of Nemp1 KO *RBCs*. **E**. Hemavet measurement of red blood cells (RBC), red blood cells distribution width (RDW) and hemoglobin (Hg) content of Nemp1KO blood from postnatal day 4 (P4) and P10. 6, 3, 5 and 4 mice were respectively analyzed for RBC, Hb and RDW measurements for P4 WT, P4 KO, P10 WT and P10 KO mice. **F**. Percentage of immature nucleated erythroblasts in peripheral blood of indicated genotypes. **G**. H&E staining and quantification of cellularity of WT and Nemp1KO BM. **H, I.** Percentage of Ter119+ (H) and Ter119+/CD71+ (I) cells in WT and Nemp1 KO BM assessed by FACS. **J.** Percentage of nucleated Ter119+/Hoechst+ cells in WT and Nemp1 KO BM measured by flow imaging. **K,L.** Percentage of Ter119+ (H) and Ter119+/CD71+ (I) cells in WT and Nemp1 KO spleen assessed by FACS. **M.** Percentage of nucleated Ter119+/Hoechst+ cells in WT and Nemp1 KO spleen measured by flow imaging. Data are shown as mean ± SD. Student’s t-test. ns. not significant, *p<0.05. **p<0.01. ***p<0.001. ****p<0.0001.

To understand the nature and origin of erythroid defects, we quantified the erythroid population of BM and spleens from adult WT and *Nemp1* KO mice by using Ter119 and CD71 markers. Compared to WT BM, *Nemp1* KO BM showed increased Ter119+ cell population (Figure 1H) that was mostly accounted for by a significantly increased population of Ter119+CD71+ erythroblasts (Figure 1I). The increase in erythroblasts was also detected by flow imaging of Hoechst+Ter119+ populations (Figure 1J). In the adult spleen, FACS and flow imaging data showed similar increases in the representation of erythroblast population (Figure 1 K, L, and M). Taken together, these data show that loss of *Nemp1* leads to a significant increase of the erythroid lineage in BM and spleens.

### Erythroid progenitors and early erythroblast populations are expanded in *Nemp1* KO mice

To trace the origin of erythroid lineage overrepresentation in *Nemp1* KO mice, we quantified hematopoietic progenitors. Long-term hematopoietic stem cells (LT-HSCs, SLAM-KSL), hematopoietic stem and progenitor cells (HSPCs, KSL, ckit+/Sca1+/Lin-) as well as common myeloid progenitors (CMP, CD34^+^CD16/32^-^Lin^-^c-Kit^+^Sca1^-^) were equally represented in WT and *Nemp1* KO BM (Figure 2 A,B,C). In contrast, the megakaryocyte-erythrocyte progenitor (MEP, CD34^-^CD16/32^-^Lin^-^c-Kit^+^Sca1^-^) population was significantly increased with a concomitant decrease of the granulocyte-macrophage progenitor (GMP, CD34^+^CD16/32^+^Lin^-^c-Kit^+^Sca1^-^) population (Figure 2C) in *Nemp1* KO BM. In agreement with this increase in MEPs, *Nemp1* KO BM cells consistently showed a higher capacity to generate BFU-E (Figure 2D). Together, these results indicate that the earliest phenotype resulting from the lack of *Nemp1* expression is the expansion of MEPs in the hematopoietic cascade.

**Figure 2:**
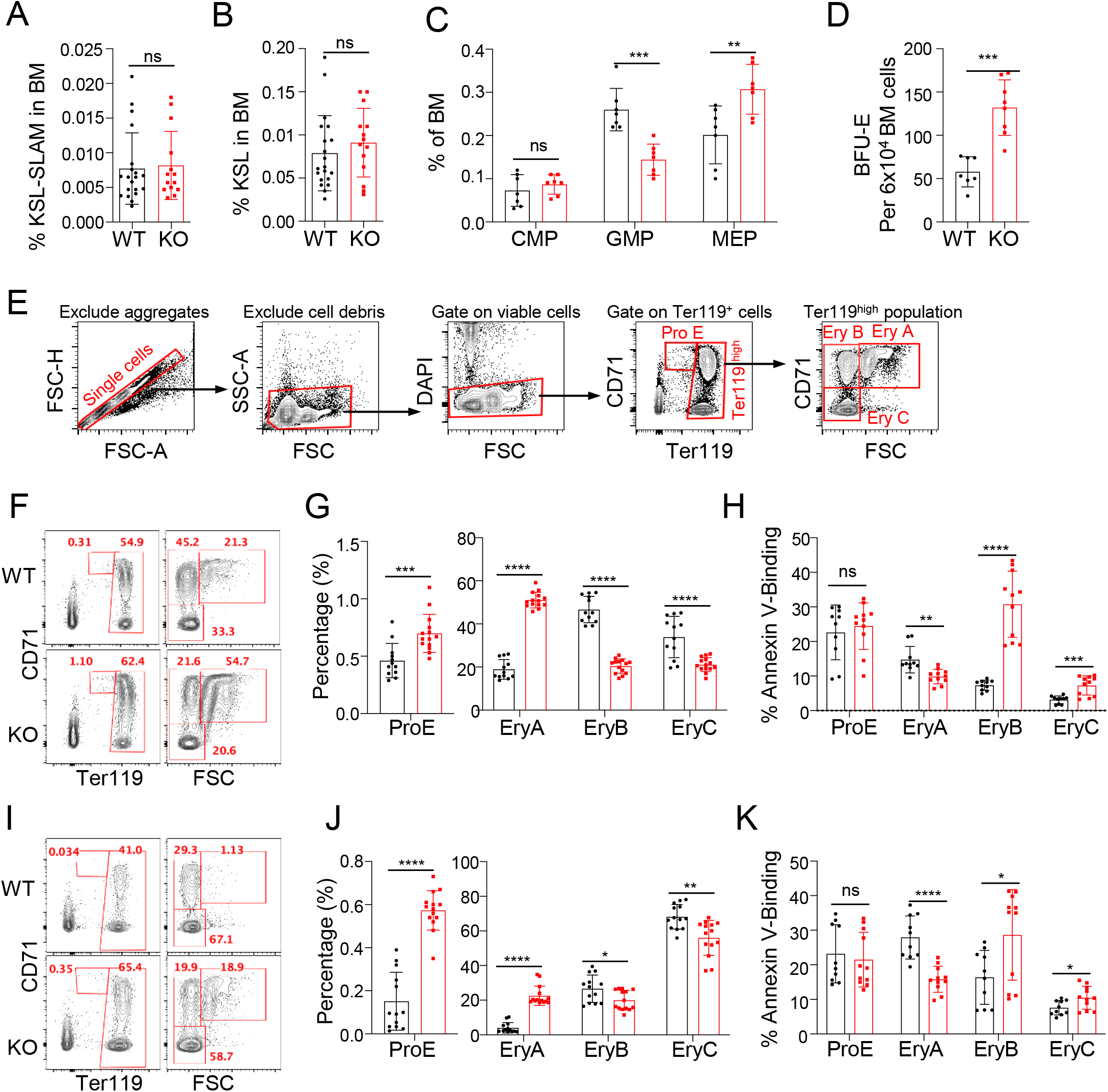
Loss of Nemp1 leads to the expansion of erythroid progenitors and early erythroblasts. Quantification of LT-HSCs (**A**), HSPCs (**B**), CMPs, GMPs and MEPs progenitors (**C**). **D.** BFU-E formation capacity of WT and Nemp1 KO BM. **E.** FACS sorting strategy used to quantify ProE and Ery A, B, C populations from whole BM and spleen. Representative FACS analyses and quantification of ProE, EryA, B and C populations in WT and Nemp1KO BM (**F, G**) and spleens (**I, J**). Apoptotic indexes of the same respective populations in WT and Nemp1 KO BM (**H**) and spleen (**K**). Data are shown as individual subject and the mean ± SD. Student’s t-test. ns. not significant, *p<0.05. **p<0.01. ***p<0.001. ****p<0.0001.

To better understand the erythroid differentiation defects in *Nemp1* KO mice, we next examined the representation of ProE, EryA, EryB, and EryC erythroblasts using the gating strategy shown in Figure 2E. Consistent with increased MEP population and higher BFU formation capacity in *Nemp1* KO BM (Figure 2C, D), the ProE and EryA populations were increased, with EryA showing a more significant increase (Figure 2F, G). By contrast, the EryB and EryC were decreased, with EryB showing a more significant decrease, in the *Nemp1* KO BM. Whereas apoptosis was mildly reduced in EryA, the apoptotic EryB population was significantly higher in *Nemp1* KO BM (Figure 2H). Similar trends were also observed in the spleen (Figure 2I, J, and K). Collectively, these data indicate that genetic ablation of *Nemp1* leads to the expansion of MEPs to EryA populations and to a decrease of EryB and EryC populations. Increased apoptosis in EryB probably accounts for the loss of RBC.

Given the compromised erythropoiesis in the BM, we suspected that spleen enlargement in *Nemp1* KO mice might be due to stress erythropoiesis. Accordingly, c-kit+/CD71+/Ter119-stress erythroid progenitors (SEPs) and Epo-responsive BFU-E and CFU-E were significantly increased in Nemp1KO spleen and splenectomy experiments also pointed to a significant contribution of splenic SEPs to ongoing erythropoiesis in Nemp1 KO mice (S2 text and S2 Fig). Collectively, these data show that genetic ablation of NEMP1 affects erythroid maturation, especially from EryA to EryB, and elicit stress erythropoiesis.

### NEMP1 is required for erythroblast NE openings, nuclear compaction and nuclear extrusion

Using a NEMP1 antibody directed towards a stretch of 15 amino acids from the C-terminal region of NEMP1 (Figure S3A) [1], NEMP1 (49.8 kDa) was detected both in lysates of sorted WT CD11b+ myeloid and Ter119+ erythroid cells but not in their KO counterparts (Figure 3A). In immunofluorescence confocal microscopy, NEMP1 was detected at the NE of ProE, EryA, B and C erythroblasts where it colocalized with LAP2, a well-established NE marker (Figure 3B). However, NEMP1 noticeably formed occasional NE puncta of higher intensity whereas LAP2 was homogenously distributed on the NE of erythroblasts (Figure 3B). NEMP1 was also detected at the NE of cKit+/Ter119-progenitors (Figure S3B). NEMP1 was undetectable at the NE of BM cells isolated from *Nemp1* KO mice demonstrating the specificity of the NEMP1 antibody (Figure S3C). Taken together, these results show that NEMP1 is ubiquitously expressed at the NE of the erythroid lineage.

**Figure 3:**
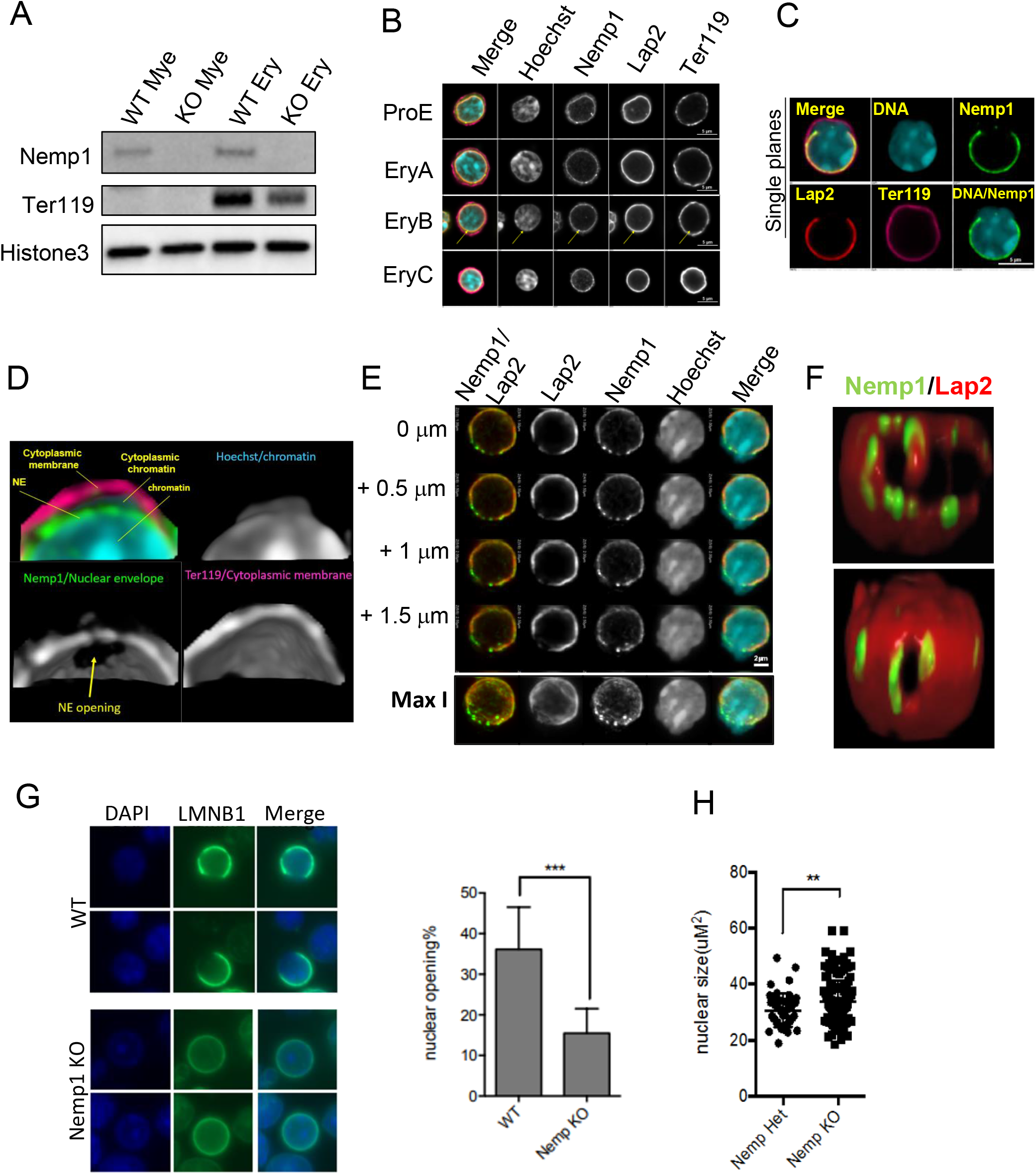
Nemp1 is required for NE openings in erythroblasts. **A.** Immunodetection of endogenous NEMP1 in lysates from WT and Nemp1 KO myeloid (CD11b+) and erythroid (Ter119+) cells. Histone3 was used as a loading control and Ter119 to confirm myeloid/erythroid sorting specificity. **B**. Nemp1 immunostaining of WT BM erythroblasts. Lap2 was used as a control for NE labeling. ProE were identified based on low Ter119 expression and low nuclear circularity. Ery A, B and C were distinguished based on increasing Ter119 intensity and decreasing nuclear diameter (7.4, 6.4 and 5.4 μm for EryA, B and C, respectively). The arrow points to discrete Nemp1 aggregates. **C.** Immunolocalization of Nemp1 and Lap2 in a WT erythroblast undergoing NE opening and chromatin release into the cytoplasm. **D.** 3D reconstruction (non-thresholded) of single confocal planes of a NE opening delineated by Nemp1 after denoising and α-blending. Note the extrusion of chromatin through the NE openings within the cytoplasm. **E.** Successive confocal planes (0.5 μm apart) of an erythroblast undergoing NE opening showing Nemp1 aggregates at NE opening sites. Note the presence of higher intensity aggregates of Nemp1 at or near NE openings delineated by Lap2. Bottom panel: Maximum intensity projection (Max I) of a Z-stack encompassing the NE opening. **F**. 3D reconstruction after α-blending of confocal slices thresholded for Nemp1 intensity showing Nemp1 aggregates at or near to NE openings. The bottom panel shows a small NE opening already decorated with Nemp1 aggregates. **G**. Confocal imaging of LaminB1 in WT and Nemp1 KO BM cells and quantification of NE opening frequencies. **H**. Quantification of nuclear area in WT and Nemp1 KO BM cells.

Late stages of erythroblast maturation are characterized by the progressive compaction of chromatin and its partial release into the cytoplasm via transient openings of the NE, which is most prominent in the polychromatophilic stage [9,10]. As shown in Figure 3C, NE openings were clearly identified with NEMP1 and LAP2 antibodies in Ter119+ erythroblasts. This phenomenon is distinct from nuclear extrusion as the cytoplasmic membrane labelled with Ter119 remains intact (Figure 3C). 3D reconstruction of confocal Z-stacks clearly emphasized NE openings delineated by NEMP1 through which chromatin protrudes into the cytoplasm (Figure 3D and S1 Video).

Interestingly, the close examination of successive confocal planes encompassing NE openings revealed the accumulation of NEMP1 into higher intensity aggregates (Figure 3E). Maximum intensity projections also showed that NEMP1 aggregates preferentially accumulated near or close to NE openings (Figure 3E, bottom panel). In contrast, the localization of LAP2 remained uniform and did not accumulate into NEMP1 aggregates. Intensity profiles showed that NEMP1 was 2 to 3 times more abundant in aggregates at NE openings by comparison to intact NE (Figure S3D). To better appreciate the spatial distribution of Nemp1 aggregates relative to NE openings in three dimensions (3D), we performed intensity thresholding of confocal slices followed by 3D reconstruction and α-blending. As shown in Figure 3F, multiple NEMP1 aggregates localized close to or at the edges of NE opening sites delineated by Lap2 (Figure 3F, top and video S2). Interestingly, small NE openings already displayed a few NEMP1 aggregates at their edges (Figure 3F, bottom panel and video S3). NEMP1 aggregates were undetectable at the NE of *Nemp1* KO erythroblasts thereby confirming the specificity of these structures (Figure S3E).

To determine if NEMP1 is required for NE openings, WT and *Nemp1* KO BM were immunostained with Lamin B1. Loss of *Nemp1* expression led to a significant decrease of NE opening frequencies suggesting a role for NEMP1 in NE openings (Figure 3G). We also observed a significant increase of nuclear size in *Nemp1* KO BM cells that may be indicative of decreased chromatin compaction (Figure 3H).

Because chromatin compaction is required for enucleation [9], we next examined whether NEMP1 is required for enucleation. As shown in Figure 4A, NEMP1 decorated the NE with occasional puncta at all described stages of enucleation [13]. We used two approaches to determine whether genetic ablation of *Nemp1* affects nuclear extrusion. First, lineage negative cells from wild type and *Nemp1* KO mice were purified from BM, cultured for 2 days in erythropoietin-containing medium and analyzed to distinguish nucleated vs non-nucleated Ter119+ cells. As shown in Figure 4B, although erythroblast differentiation was not affected, the ratio of nucleated erythroblasts (DNA+/Ter119+) vs RBC (DNA-/Ter119+) was increased in cultures derived from *Nemp1* KO BM. In agreement with these data and using flow imaging as a second approach, we consistently measured a higher ratio of nucleated erythroblasts (Hoechst+/Ter119+) vs RBC (DNA-/Ter119+) in *Nemp1* KO BM by comparison to WT BM cells (Figure 4C). Taken together, these results indicate that NEMP1 plays a role in NE openings and nuclear extrusion.

**Figure 4:**
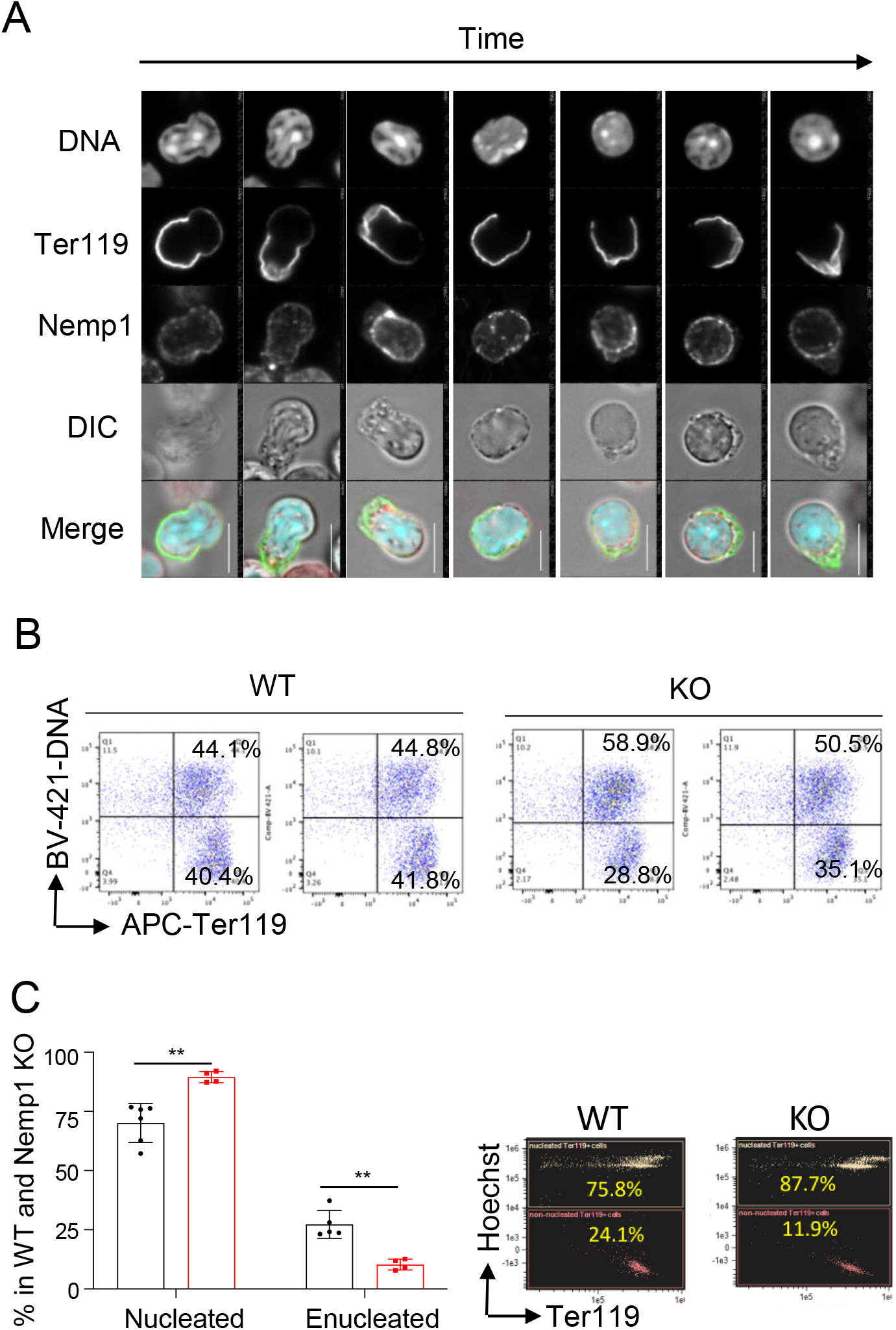
Nemp1 is required for erythroblasts enucleation. **A.** Immunolocalization of Nemp1 at different stages of enucleation. Scale bars: 5 μm. **B.** DNA and Ter119 FACS analysis of lineage-negative cells purified from WT and KO BM and subsequently cultured in erythropoietin-containing medium for 2 days. Note the higher nucleated/enucleated ratio in Nemp1 KO cells. **C**. Quantification of nucleated and enucleated Ter119+ cells in WT and Nemp1 KO BM measured by flow imaging. Right: representative quantification graphs of nucleated (Hoechst+Ter119+) and enucleated (Hoechst-Ter119+) in WT and Nemp1KO BM in flow imaging.

## DISCUSSION

In this work, we show that loss of the integral transmembrane NE protein NEMP1 results in erythropoietic defects consisting of the expansion of the erythroid lineage in BM and spleens and erythroid maturation defects during terminal erythropoiesis (Figure 5). As a consequence, *Nemp1* KO mice display anemia in neonates and splenomegaly associated with stress erythropoiesis in adult mice.

**Figure 5:**
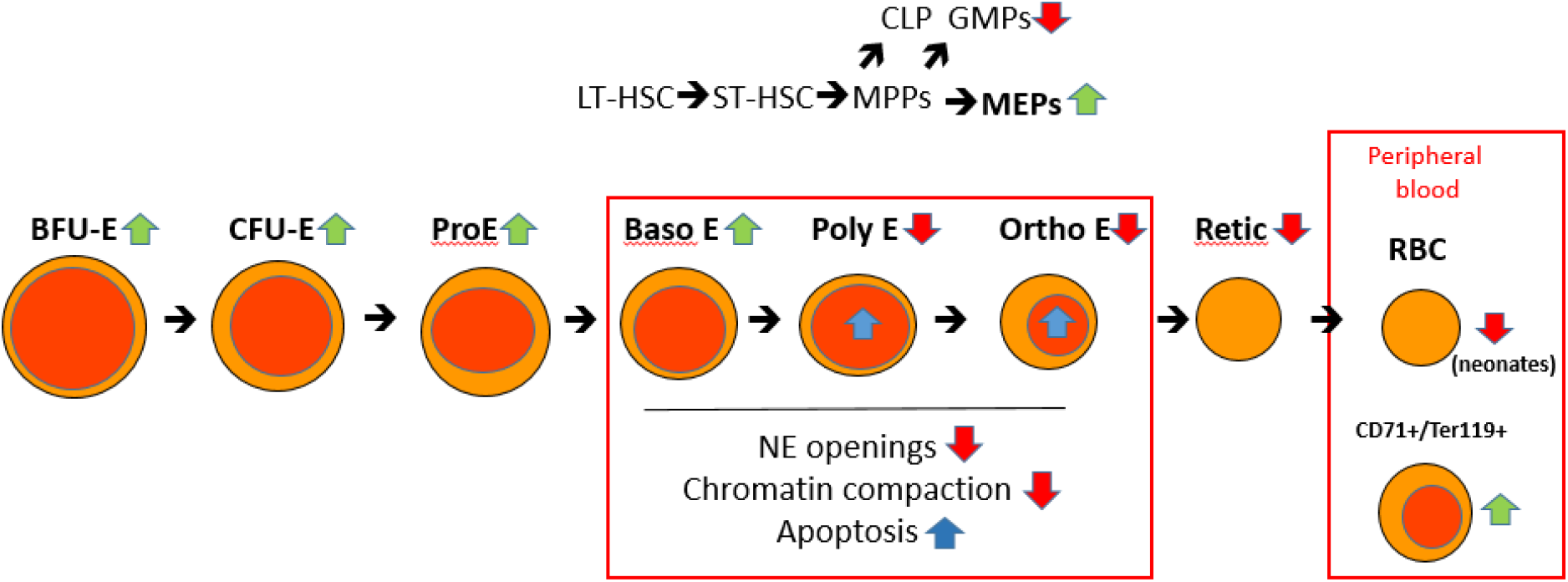
Graphical depiction of Nemp1 phenotypes and functions during erythropoiesis. Green and red arrows respectively depict increased or decreased populations in Nemp1KO by comparison to WT cells or biological processes affected by the lack of expression of NEMP1. Blue arrows depict increased apoptosis. See text for details.

The higher capacity of *Nemp1* KO BM and spleens to generate BFU-E and CFU-E and the significant expansion of ProE and EryA populations together show that erythroid expansion is taking place throughout the EryA stage in *Nemp1* KO mice. MEP expansion was notably associated with a significant decrease in GMPs, suggesting that *Nemp1* may have an additional biological function in non-erythroid lineages. Alternatively, GMP decrease may reflect compensatory mechanisms in response to erythroid lineage expansion.

We speculate that the overrepresentation of erythroid progenitors and EryA as well as the stress erythropoiesis we observed in spleen originate from a feedback loop due to the massive apoptotic loss of EryB erythroblasts. Indeed, in contrast to their progenitors and EryA precursors, EryB and EryC populations were markedly decreased in *Nemp1* KO BM and RBC counts were significantly lower in the peripheral blood of neonate and young *Nemp1* KO mice. This decrease of EryB and EryC populations was accompanied by a marked increase of apoptosis that was especially pronounced in EryB from BM. We reason that this apoptosis stems from key biological functions of NEMP1 in NE openings during terminal erythropoiesis. First, *Nemp1* transcripts are significantly upregulated during terminal differentiation [14] and *Nemp1* expression level peaks in EryB/polychromatophilic erythroblasts [15]. Second, in erythroblasts undergoing NE openings, NEMP1 accumulates into aggregates that preferentially localized near or at NE openings. Importantly, this aggregation is specific to NEMP1 as LAP2 remains uniformly distributed at NE openings. It is possible that increased levels of NEMP1 expression reported in proteomic screens at that differentiation stage [15] reflects the accumulation of Nemp1 into aggregates. Finally, we directly show that lack of *Nemp1* expression in erythroblasts is linked to reduced frequencies of NE openings. Taken together, we propose that the increased apoptosis measured in *Nemp1* KO EryB and EryC stems from the requirement of NEMP1 aggregates for efficient NE openings. In support of this idea, inhibition of NE openings is directly linked to induction of cell death in G1ER cells [9].

An alternative but not exclusive origin of elevated apoptosis in EryB erythroblasts may also stem from a biological function of NEMP1 in chromatin organization. Indeed, we recently reported the interaction of NEMP1 with LEM domain-containing proteins EMERIN, LAP2 and MAN1 [1] that are also expressed in erythroblasts [15] and play essential roles in chromatin organization in cooperation with BAF [16]. In addition, the genetic ablation of NEMP proteins in plants (PNET2) leads to major disruptions of higher-order chromatin organization [4] and *Nemp1* KO erythroblasts displayed larger nuclear sizes. To that regard, it is interesting to note that the genetic inactivation of LAP2α, a LAP2 isoform devoid of transmembrane domain and involved in the stabilization of higher order chromatin [17], also leads to the overrepresentation of the erythroid lineage [18]. Impaired nuclear extrusion in *Nemp1* KO erythroblasts may also contribute to increased apoptosis levels but future studies are needed to determine whether NEMP1 intrinsically affects nuclear extrusion or if decreased enucleation in *Nemp1* KO BM is a mere consequence of impaired NE openings.

We detected the expression of NEMP1 at the NE of erythroblasts and cKit+ progenitors by immunofluorescence microscopy and in purified Ter119+ erythroid cells by immunoblotting which is in agreement with its identification in proteomic analyses of the human erythroid cells[15]. By contrast, the same proteomic analyses show that human erythroblasts do not express *Nemp2,* a homolog of *Nemp1* [15]. Accordingly, *Nemp2* KO mice do not display any splenomegaly or blood defects, and *Nemp1* phenotypes were not worsened in *Nemp1/2* double knockout (DKO) mice (data not shown). Taken together, we conclude that *Nemp1* is specifically required for the normal homeostasis of the erythroid lineage.

The precise function of NEMP1 aggregates at NE openings requires more investigation. Biochemical studies showing that Nemp1 oligomerizes via its transmembrane domains [3] suggest that NEMP1 may aggregate through multimerization upon increased expression levels detected in transcriptomic and proteomic studies [14,15]. Because we know that NEMP1 supports NE stiffness in the germline and cultured cells[1], it is possible that Nemp1 aggregates mechanically support the nuclear envelope during the physical stresses of NE openings.

To our knowledge, this is the first report describing the requirement for an integral transmembrane protein of the NE in erythropoiesis. Indeed, pathologies linked to mutations of NE proteins or the nuclear lamina, globally termed “nuclear envelopathies” and “laminopathies”, have not previously been associated to erythropoietic defects[19,20]. Similarly to the involvement of multiple NE proteins in specific pathologies, Nemp1 deficiency specifically underlies female infertility and erythropoietic defects in mice. This further fuels the concept of tissue-specific composition of NE proteins whose mutations results in tissue-specific pathologies[21].

Repetitive NE openings during terminal differentiation provides a relatively little-known yet powerful opportunity to study NE dynamics and remodeling *in vivo*. To that regard, Zhao et al.[7,9] have shown that NE openings are blocked by inhibition of caspase-3 or through the expression of a caspase-3 non-cleavable Lamin B1 mutant. As a result, histone release from the nucleus, chromatin condensation and the terminal differentiation of erythroid cells are also affected *in vitro.* These data and our current findings therefore further stress the biological relevance of NE openings during terminal erythropoiesis. Finally, because Caspase-3 KO mice show relatively mild erythroid defects most likely due to *in vivo* compensatory pathways[9], *Nemp1* KO mice provide the first mouse model of acute erythropoietic defects linked to NE openings deficiency. In conclusion, our results uncovered the involvement of *Nemp1* in NE openings and enucleation in erythroblasts and its requirement for normal erythropoiesis.

## MATERIAL AND METHODS

### Animals

Animal protocols used in this study strictly adhered to the ethical and sensitive care and use of animals in research and were approved by the Washington University School of Medicine Animal Studies Committee (Animal Welfare Insurance Permit #A-3381-01, protocol#21-0206). *mNemp1* (Nemp1^em# (TCP) McNeill^) CRISPR KO allele was obtained by CRISPR-Cas9–mediated deletion of exon3 that is present in all *mNemp1* transcripts (Toronto Center for Phenogenomics). Mice were generated and maintained on a C57Bl6N background [1].

### Antibodies

Rat Ter119-Alexa647 (Biolegend, #116218, 1:200), rabbit Nemp1 (1:1,000), mouse Lap2 (BD Biosciences, #611000, 1:200), Rat ckit-Alexa647 (#105817, 1:200), goat LaminB1 (Santa Cruz Biotechnology) were used for immunofluorescence microscopy. Fluotag-X4 anti-Rabbit IgG-Atto488 (Nanotag, #N2404, 1:500) and FluotagX2 anti-mouse IgG-AberioSTAR-580 (Nanotag, #N1202, 1:500) were used as secondary antibodies. Rat Ter-119-PE (Biolegend, #116208, 1:200), Rat CD71-APC (Biolegend, #113820, 1:200) were used for FACS and flow imaging. Histone3 (Abcam, #1791, 1:2000), Nemp1 (1:1000) and Ter119-Biotin (Biolegend, #116204, 1:500) were used for immunoblotting.

### Bone marrow and spleen cells collection

Two to four month-old mice were euthanized by CO_2_ inhalation. Legs were separated at the pelvic-hip joint and femurs and tibia cleaned off from tendons and muscle tissues in cold PBS. Bones were cut on their extremities and transferred into collection units consisting of 0.5 ml tubes with pre-perforated (18 gauge needle) bottoms inserted in 1.5 ml collection tubes. Collections units were centrifuged 4 min at 6000 RPM at 4C. Pellets of BM cells were then resuspended in 1 ml of FACS buffer (0.5% BSA and 2mM EDTA in PBS), transferred to a 40uM cell strainer and washed with 10 ml of FACS buffer. Cell suspensions were then spinned down for 3 min at 3,000 RPM. Pellets were resuspended in 2 ml of FACS buffer and fixed by adding 3.5 ml of fixation/permeabilization buffer (BD Biosciences) and rocking overnight at 4C. Cells were then washed in permeabilization buffer (BD Biosciences) through three cycles of centrifugation for 3 min at 850g and stored at 4C for further use.

### Flow imaging

One million fixed BM and spleen cells were immunolabelled for one hour at room temperature with Ter119-PE (1:200, Biolegend) and 1ug/ml Hoechst 33342 (Thermofisher) in permeabilization buffer and then washed in PBS through three cycles of centrifugation for 3 min at 850g. The final pellet was resuspended in 40 μl PBS for image flow analysis. Data were acquired on an AMNIS ImageStreamX multispectral imaging flow cytometer (Luminex) using the Inspire software package. All images were acquired with the 60X objective with Hoechst (405 laser line), Ter119PE (488 laser line) and brightfield imaged in channels 1, 3 and 4, respectively. Laser intensities were adjusted to avoid signal saturation. Single fluorophore labeling were used to build a compensation matrix. Post-acquisition data analyses were performed with the IDEAS software package. For measurements of Ter119+ erythroid populations, cells in focus (gradient RMS) were gated on Hoechst+ cells that were subsequently plotted for Ter119 intensity.

### Immunofluorescence confocal microscopy

For immunostaining of intracellular epitopes, fixed BM cells were washed and permeabilized three times in Perm/wash buffer (BD Biosciences) supplemented with 0.1% TritonX100 and further incubated in the same buffer with primary antibodies overnight at 4C. After three washes, cells were incubated with secondary antibodies and with fluorescently labeled antibodies against extracellular epitopes for 2 hours at room temperature and then washed three times with three cycles of centrifugation for 3 min at 850g. Cells pellet were resuspended in 20ul of PBS. 3 μl of cell suspension were mixed with 10ul of fluorescence mounting medium (Dako) and mounted for downstream confocal imaging. All images were acquired on a Nikon confocal microscope with a 1.4 NA 100X objective. Images denoising and 3D reconstruction were performed with the NIS-Element software package suite.

### Analysis of mouse peripheral blood

Whole blood was collected by venipuncture of the facial vein and immediately transferred in blood collection tubes (BD Microtainer). Blood samples were mixed and placed under the Hemavet HV950 probe (Drew Scientific, Inc.) for analysis using reagents from the LV-PAK (Drew Scientific Inc.). Multi-Trol mouse serum controls (Drew Scientific, Inc.) were used for calibration of the Hemavet HV950. Collected blood was also spread on glass slides for Wright-Giemsa staining according to the manufacturer’s protocol (Wright-Giemsa Stain Modified, Sigma-Aldrich).

### In vitro colony-forming assay

Methylcellulose colony-forming assay were performed using Epo-only MethoCult 3334 (Stem Cell Technologies) according to the manufacturer’s instructions. BM (6×10^4^ cells/mL) or spleen (1×10^5^ cells/mL) cells were mixed with M3334 methylcellulose and plated in triplicates using 35mm Petri-dishes. Cultures were maintained in a humidified incubator at 37°C, 5% CO_2_. CFU-E colonies were counted after 2-3 days of culture. BFU-E colonies were counted 5 days after culturing.

### Acute Anemia and Splenectomy

Acute hemolytic anemia was induced by intraperitoneal injection of phenylhydrazine (PHZ) (Sigma) with a single dose of 100mg/kg. Splenectomy was performed at the Hope Center for Neurological Disorder at Washington University in St. Louis. Approximately 1 month later, the splenectomized mice were induced by PHZ for hemolytic anemia. Blood samples were collected by venipuncture of the facial vein at different time point, and hematologic parameters measured on a HemaVet HV950 complete blood count instrument.

### Flow cytometry and cell sorting

BM and spleen cells from WT or Nemp1 KO mice were dissociated, resuspended in PBS/0.5% BSA and passed through a 40uM cell strainer to obtain single-cell suspension before antibody staining. Analysis of erythroid maturation using CD71 and Ter119 was conducted as previously described [22]. Freshly isolated cells were stained at 4°C in PBS/0.5% BSA with purified anti-mouse CD16/32 to block Fc receptors and incubated with PE-Ter119 (TER-119) and APC-CD71 (R17217) antibodies for 30 minutes at 4°C. DAPI was used to exclude dead cells from analysis. Where apoptosis was measured, immunostaining for Ter119 and CD71 was followed by a 15-minute incubation with FITC-conjugated Annexin-V and propidium iodide following the manufacturer’s protocol (BD Biosciences). Flow cytometry was carried out on BD LSR II machine and BD FACSAria II was used for cell sorting. Gate strategy was performed as previously described [22]. The ProE gate contains CD71^high^Ter119^intermediate^. The Ter119^high^ cells are further analyzed. Here CD71^high^ cells are subdivided into less mature, large EryA erythroblasts (CD71^high^Ter119^high^FSC^high^) and smaller, more mature EryB erythroblasts (CD71^high^Ter119^high^FSC^low^). The most mature erythroblasts subset is EryC (CD71^low^Ter119^high^FSC^low^).

HSPC and committed progenitors staining were conducted as described previously [23]. Cells from BM were harvested in PBS/0.5% BSA, and quickly lysed with RBC lysis buffer for 1 min at 4°C. Cells were then stained with PE-Cy7 conjugated anti-Gr-1 (RB6-8C5), -Mac1 (M1/70), -B220 (RA3-6B2), -Ter119 (TER-119), -CD3 (17A2), in combination with APC-e780-c-Kit (2B8), PerCP-Cy5.5-Sca1 (D7), APC-CD48 (HM48-1), PE-CD150 (TC-12F12.2), FITC-CD34 (RAM34) and BV421-CD16/32 (93) antibodies for 30min on ice. Flow cytometry was carried out on BD Symphony A3 machine. Data were analyzed on FlowJo software (FlowJo, LLC). Different committed progenitors were defined as granulocyte-monocyte progenitor (GMP, CD34^+^CD16/32^+^Lin^-^c-Kit^+^Sca1^-^) cells, common myeloid progenitor (CMP, CD34^+^CD16/32^-^ Lin^-^c-Kit^+^Sca1^-^) cells and megakaryocyte-erythrocyte progenitor (MEP, CD34^-^CD16/32^-^ Lin^-^c-Kit^+^Sca1^-^).

## ACKNOWLEDGMENTS

The authors would like to thank the Centre for Phenogenomics (TCP), Sinai Health System, Toronto, Ontario, for assistance with blood analysis and the Mouse Genetics Core at Washington University School of Medicine in St Louis (Dr. Mia Wallace) for mouse husbandry and genotyping.

## SUPPLEMENTAL FIGURE CAPTIONS

***S1 Fig: Nemp1 KO mice have peripheral blood defects:** Hemavet measurement of red blood cells (RBC), red blood cells distribution width (RDW) and hemoglobin (Hg) content of Nemp1KO and WT blood from P4 to P150. N: number of biological replicates for each time point and genotype. Data are shown as mean ± SD. Student’s t-test. ns. not significant, *p<0.05. **p<0.01. ***p<0.001. ****p<0.0001*.

***S2 Fig: Nemp1 KO mice display increased splenic stress erythropoiesis:** Representative FACS analysis (**A**) and quantification (**B**) of SEPs in dissociated WT and Nemp1 KO spleens. Comparisons of BFU-E (**C**) and CFU-E (**D**) formation capacity between WT and Nemp1KO dissociated spleens. Data are shown as individual subject and the mean* ± SD. Student’s t-test. ****p<0.0001. *Complete blood count (CBC) analysis of WT (n=5) and KO (n=5) mice during the entire recovery period following PHZ treatment (100mg/kg). Data are show as geometric mean* ± 95% of difference. Using two-way ANOVA test: *, p=0.0112. *(**E**). CBC analysis of WT (n=6) and KO (n=6) mice after splenectomy. Data are show as geometric mean* ± 95% of difference. Using two-way ANOVA test: **, p=0.0012 for RBC comparison; p=0.0022 for Hb comparison; p=0.0045 for MCV comparison; ***, p=0.0007 *(**F**). CBC analysis of WT (n=6) and KO (n=6) mice after splenectomy plus PHZ treatment (100mg/kg). Data are show as geometric mean* ± 95% of difference. Using two-way ANOVA test: *, p=0.0102; **, p=0.0078 for RBC comparison; p=0.0013 for Hb comparison; ****, p < 0.0001; *(**G**)*.

***S3 Fig: Nemp1 localizes at the NE of hematopoietic cells and aggregates at NE openings**. **A**. Topology of NEMP1 at the NE with the Nemp1 antibody epitope denoted in green. INM, ONM: inner and outer nuclear membrane, respectively. **B.** Immunolocalization of endogenous Nemp1 in cKit+ progenitors by confocal microscopy (top panel). A Ckit-/Ter119low ProE is shown for comparison (Bottom panel). **C**. Intensity profile lines showing Nemp1 peaks (green, nuclear envelope) that are distinct from Ter119 peaks (cytoplasmic membrane) in WT BM. Nemp1 peaks are not present in Nemp1 KO BM. **D.** Intensity profile lines comparing the intensity of Nemp1 in aggregates at the edge of NE openings to the intensity of Nemp1 in intact NE. Note the lack of accumulation of either Lap2 (red trace) or chromatin (blue trace) in Nemp1 aggregates. Right panel: Maximum intensity projection (Max I) of the same cell showing the preferential accumulation of Nemp1aggregates near or at the edge of the NE opening. **E.** Maximum intensity projection of WT (top) or Nemp1 KO (bottom) erythroblasts undergoing NE opening. Note the absence of Nemp1 aggregates in Nemp1KO erythroblasts thereby confirming their specificity*.

***S1 Video:** Non-tresholded 3D reconstruction from confocal planes of an erythroblast undergoing nuclear envelope opening*

***S2, S3 Videos**: 3D reconstruction from confocal planes (thresholded for Nemp1 intensity to emphasize Nemp1 aggregates, top panels) of erythroblasts undergoing a large (top) or small (bottom) nuclear envelope opening. Note the presence of Nemp1 aggregates near or close to NE openings*.

