## Supplemental Video S1-3 for "The inner nuclear membrane protein NEMP1 is required for nuclear envelope openings and enucleation of erythroblasts during erythropoiesis"

### Slide 1
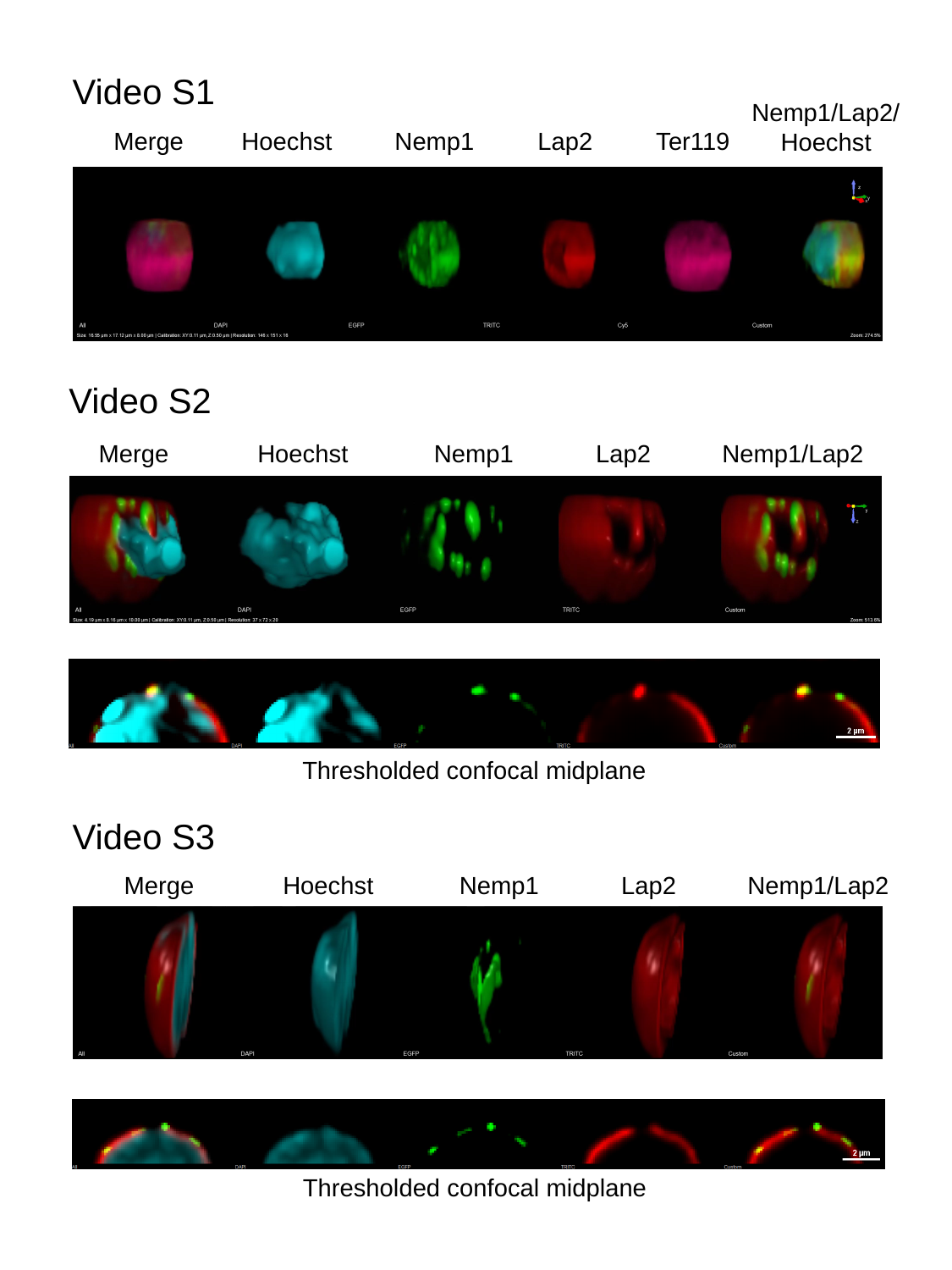

Video S1
Nemp1/Lap2/
Hoechst
Merge
Hoechst
Nemp1
Lap2
Ter119
Video S2
Merge
Hoechst
Nemp1
Lap2
Nemp1/Lap2
Thresholded confocal midplane
Video S3
Merge
Hoechst
Nemp1
Lap2
Nemp1/Lap2
Thresholded confocal midplane
