## Supplemental Figures S1-3 for "The inner nuclear membrane protein NEMP1 is required for nuclear envelope openings and enucleation of erythroblasts during erythropoiesis"

### Slide 1
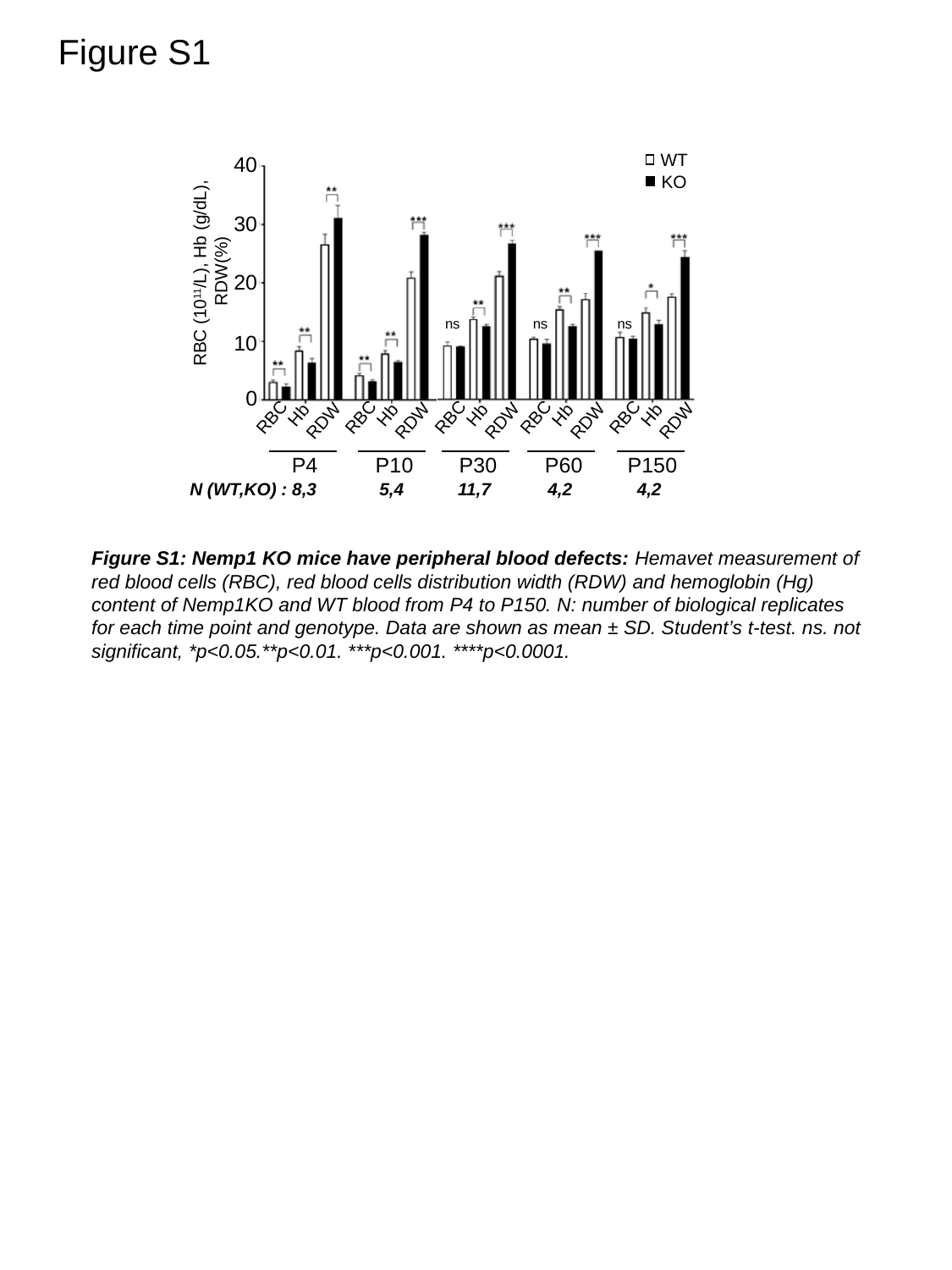

Figure S1
WT
KO
40
30
RBC (10^11^/L), Hb (g/dL),
 RDW(%)
20
10
0
Hb
Hb
Hb
Hb
Hb
RBC
RBC
RBC
RBC
RBC
RDW
RDW
RDW
RDW
RDW
P4
P10
P30
P60
P150
ns
ns
ns
N (WT,KO) : 8,3
5,4
11,7
4,2
4,2
Figure S1: Nemp1 KO mice have peripheral blood defects: Hemavet measurement of red blood cells (RBC), red blood cells distribution width (RDW) and hemoglobin (Hg) content of Nemp1KO and WT blood from P4 to P150. N: number of biological replicates for each time point and genotype. Data are shown as mean ± SD. Student’s t-test. ns. not significant, *p<0.05.**p<0.01. ***p<0.001. ****p<0.0001.

### Slide 2
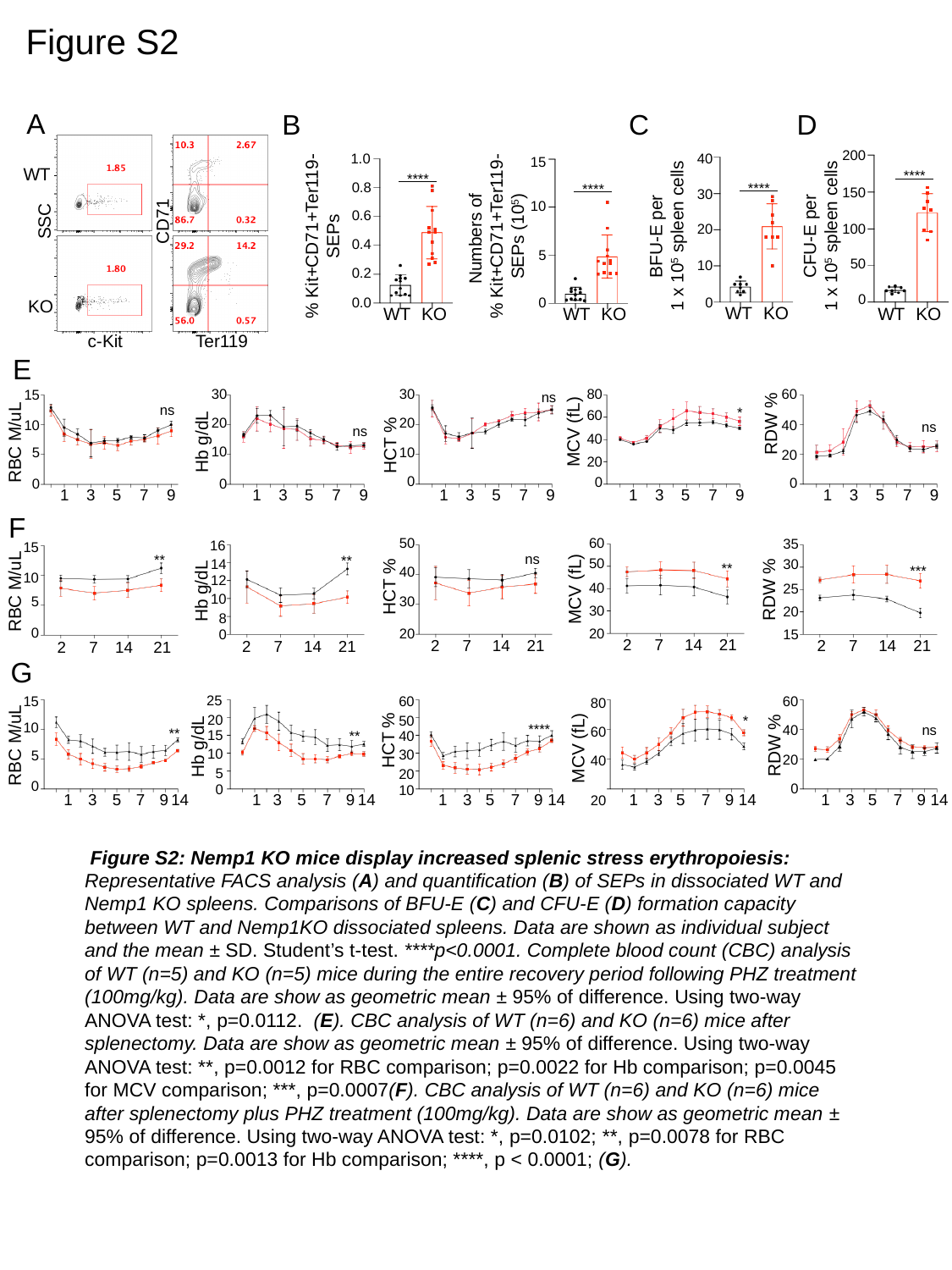

Figure S2
A
B
C
D
200
40
1.0
15
WT
****
****
0.8
****
****
150
30
10
Numbers of
% Kit+CD71+Ter119-
SEPs (10^5^)
0.6
SSC
CD71
% Kit+CD71+Ter119-
SEPs
BFU-E per
1 x 10^5^ spleen cells
CFU-E per
1 x 10^5^ spleen cells
20
100
0.4
5
50
10
0.2
0
0.0
0
0
KO
WT
KO
WT
KO
WT
KO
WT
KO
c-Kit
Ter119
E
80
60
30
30
15
ns
ns
*
60
RDW %
20
20
10
40
ns
MCV (fL)
ns
40
Hb g/dL
RBC M/uL
HCT %
10
10
5
20
20
0
0
0
0
0
1
3
5
7
9
1
3
5
7
9
1
3
5
7
9
1
3
5
7
9
1
3
5
7
9
F
60
50
35
16
15
ns
**
**
50
30
14
**
***
40
10
12
HCT %
MCV (fL)
40
RDW %
25
Hb g/dL
RBC M/uL
10
30
5
30
20
8
0
20
20
0
15
2
7
14
21
2
7
14
21
2
7
14
21
2
7
14
21
2
7
14
21
G
25
15
60
60
80
20
50
*
10
****
40
ns
60
**
15
**
40
HCT %
RBC M/uL
RDW %
Hb g/dL
MCV (fL)
10
30
5
20
40
5
20
0
0
0
10
1
3
5
7
9
14
1
3
5
7
9
14
1
3
5
7
9
14
1
3
5
7
9
14
1
3
5
7
9
14
20
 Figure S2: Nemp1 KO mice display increased splenic stress erythropoiesis: Representative FACS analysis (A) and quantification (B) of SEPs in dissociated WT and Nemp1 KO spleens. Comparisons of BFU-E (C) and CFU-E (D) formation capacity between WT and Nemp1KO dissociated spleens. Data are shown as individual subject and the mean ± SD. Student’s t-test. ****p<0.0001. Complete blood count (CBC) analysis of WT (n=5) and KO (n=5) mice during the entire recovery period following PHZ treatment (100mg/kg). Data are show as geometric mean ± 95% of difference. Using two-way ANOVA test: *, p=0.0112. (E). CBC analysis of WT (n=6) and KO (n=6) mice after splenectomy. Data are show as geometric mean ± 95% of difference. Using two-way ANOVA test: **, p=0.0012 for RBC comparison; p=0.0022 for Hb comparison; p=0.0045 for MCV comparison; ***, p=0.0007(F). CBC analysis of WT (n=6) and KO (n=6) mice after splenectomy plus PHZ treatment (100mg/kg). Data are show as geometric mean ± 95% of difference. Using two-way ANOVA test: *, p=0.0102; **, p=0.0078 for RBC comparison; p=0.0013 for Hb comparison; ****, p < 0.0001; (G).

### Slide 3
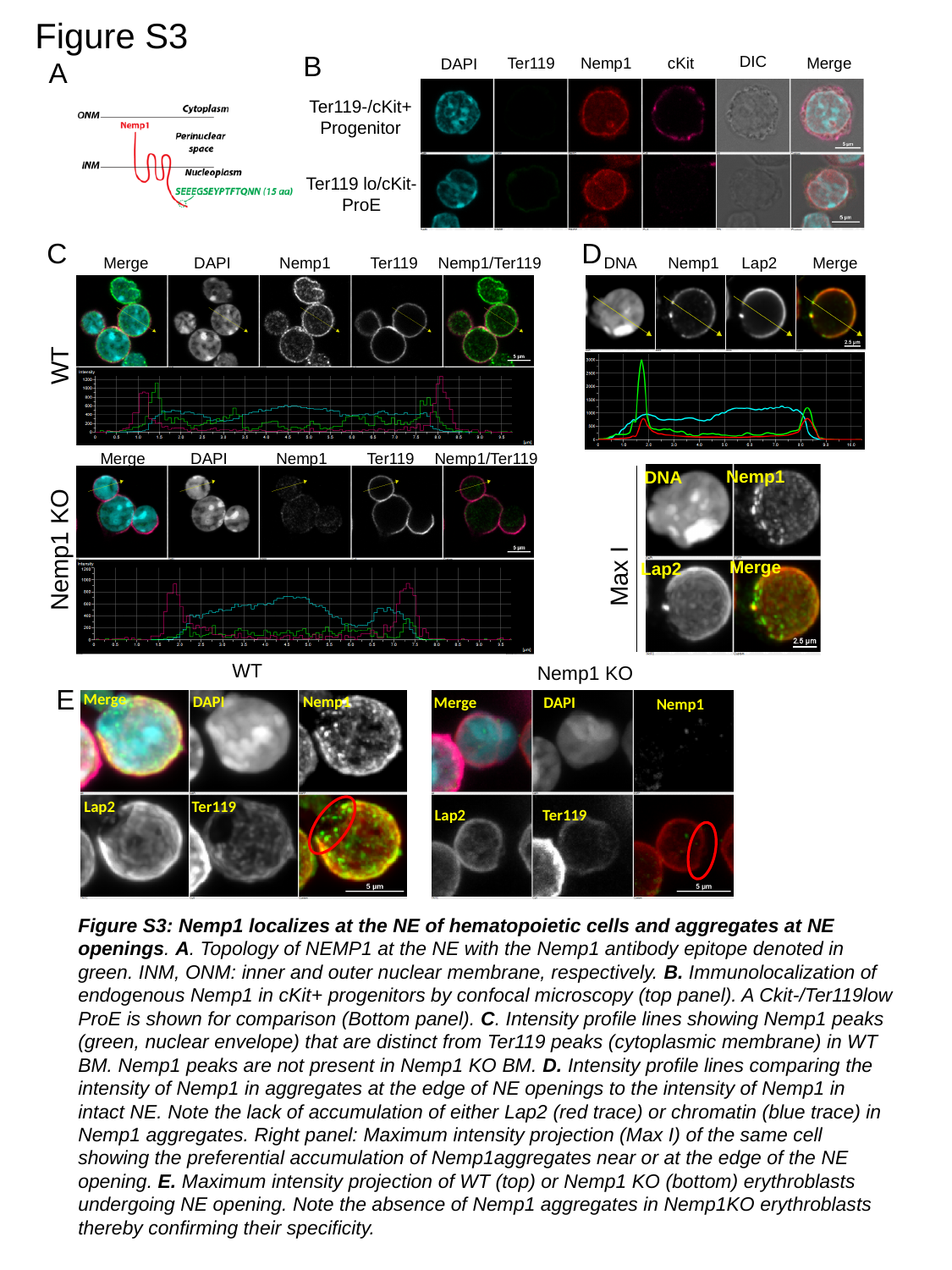

Figure S3
B
DIC
Ter119
Nemp1
cKit
Merge
DAPI
A
Ter119-/cKit+
Progenitor
Ter119 lo/cKit-
ProE
C
D
Merge
DAPI
Nemp1
Ter119
Nemp1/Ter119
WT
Merge
DAPI
Nemp1
Ter119
Nemp1/Ter119
Nemp1 KO
DNA
Nemp1
Lap2
Merge
Nemp1
DNA
Merge
Lap2
Max I
WT
Nemp1 KO
Merge
DAPI
Nemp1
Lap2
Ter119
Merge
DAPI
Nemp1
Lap2
Ter119
E
Figure S3: Nemp1 localizes at the NE of hematopoietic cells and aggregates at NE openings. A. Topology of NEMP1 at the NE with the Nemp1 antibody epitope denoted in green. INM, ONM: inner and outer nuclear membrane, respectively. B. Immunolocalization of endogenous Nemp1 in cKit+ progenitors by confocal microscopy (top panel). A Ckit-/Ter119low ProE is shown for comparison (Bottom panel). C. Intensity profile lines showing Nemp1 peaks (green, nuclear envelope) that are distinct from Ter119 peaks (cytoplasmic membrane) in WT BM. Nemp1 peaks are not present in Nemp1 KO BM. D. Intensity profile lines comparing the intensity of Nemp1 in aggregates at the edge of NE openings to the intensity of Nemp1 in intact NE. Note the lack of accumulation of either Lap2 (red trace) or chromatin (blue trace) in Nemp1 aggregates. Right panel: Maximum intensity projection (Max I) of the same cell showing the preferential accumulation of Nemp1aggregates near or at the edge of the NE opening. E. Maximum intensity projection of WT (top) or Nemp1 KO (bottom) erythroblasts undergoing NE opening. Note the absence of Nemp1 aggregates in Nemp1KO erythroblasts thereby confirming their specificity.
