## Supplemental figure legends S1-3 for "The inner nuclear membrane protein NEMP1 is required for nuclear envelope openings and enucleation of erythroblasts during erythropoiesis"

***Figure S1: Nemp1 KO mice have peripheral blood defects:*** *Hemavet measurement of red blood cells (RBC), red blood cells distribution width (RDW) and hemoglobin (Hg) content of Nemp1KO and WT blood from P4 to P150. N: number of biological replicates for each time point and genotype. Data are shown as mean ± SD. Student’s t-test. ns. not significant, *p<0.05.**p<0.01. ***p<0.001. ****p<0.0001.*

***Figure S2: Nemp1 KO mice display increased splenic stress erythropoiesis:*** *Representative FACS analysis (****A****) and quantification (****B****) of SEPs in dissociated WT and Nemp1 KO spleens. Comparisons of BFU-E (****C****) and CFU-E (****D****) formation capacity between WT and Nemp1KO dissociated spleens. Data are shown as individual subject and the mean* ± SD. Student’s t-test. *****p<0.0001. Complete blood count (CBC) analysis of WT (n=5) and KO (n=5) mice during the entire recovery period following PHZ treatment (100mg/kg). Data are show as geometric mean* ± 95% of difference. Using two-way ANOVA test: *, p=0.0112. *(****E****). CBC analysis of WT (n=6) and KO (n=6) mice after splenectomy. Data are show as geometric mean* ± 95% of difference. Using two-way ANOVA test: **, p=0.0012 for RBC comparison; p=0.0022 for Hb comparison; p=0.0045 for MCV comparison; ***, p=0.0007*(****F****). CBC analysis of WT (n=6) and KO (n=6) mice after splenectomy plus PHZ treatment (100mg/kg). Data are show as geometric mean* ± 95% of difference. Using two-way ANOVA test: *, p=0.0102; **, p=0.0078 for RBC comparison; p=0.0013 for Hb comparison; ****, p < 0.0001; *(****G****).*

***Nemp1* KO mice display increased splenic stress erythropoiesis in homeostatic conditions**

Given the compromised erythropoiesis in the BM, we assessed whether spleen enlargement in *Nemp1* KO mice might be due to stress erythropoiesis. Previous studies have shown that the splenic cKit+CD71med/-Ter119lo/- cell population contains stress BFU-Es^15^. When WT and *Nemp1* KO cKit+ cells from the spleen were assessed for CD71 and Ter119 expression, *Nemp1* KO spleens showed a ~5-fold increase in c-kit+/CD71+/Ter119- erythroid progenitors compared to WT spleens (Figure 3A,B). Consistently, Epo responsive BFU-E and CFU-E were significantly increased in *Nemp1* KO spleens (Figure 3C, D). If indeed *Nemp1* KO mice have a higher number of stress-erythroid progenitors (SEP), we expected that *Nemp1* KO mice might recover relatively normally from acute anemia induced by phenylhydrazine (PHZ). Thus, WT and *Nemp1* KO mice were treated with PHZ and monitored for peripheral RBC recovery. As expected, *Nemp1* KO mice recovered similarly to WT mice from PHZ induced anemia (Figure 3E). We monitored peripheral RBCs after splenectomy to assess the splenic SEP contribution to ongoing erythropoiesis. We found that RBCs, Hb, and HCTs in the periphery were greatly reduced in *Nemp1* KO mice compared to WT mice (Figure 3F). Mean corpuscular volume (MCV) and RDW measures were significantly increased in *Nemp1* KO mice (Figure 3F). When splenectomized WT and *Nemp1* KO mice were challenged with PHZ, *Nemp1* KO mice showed much reduced recovery of RBC, Hb, and HCT (Figure 3G). Collectively, these data suggest that *Nemp1* deficient mice have increased stress erythropoiesis occurring in the spleen.

***Figure S3: Nemp1 localizes at the NE of hematopoietic cells and aggregates at NE openings****.* ***A****. Topology of NEMP1 at the NE with the Nemp1 antibody epitope denoted in green. INM, ONM: inner and outer nuclear membrane, respectively.* ***B.*** *Immunolocalization of endog*e*nous Nemp1 in cKit+ progenitors by confocal microscopy (top panel). A Ckit-/Ter119low ProE is shown for comparison (Bottom panel).* ***C****. Intensity profile lines showing Nemp1 peaks (green, nuclear envelope) that are distinct from Ter119 peaks (cytoplasmic membrane) in WT BM. Nemp1 peaks are not present in Nemp1 KO BM.* ***D.*** *Intensity profile lines comparing the intensity of Nemp1 in aggregates at the edge of NE openings to the intensity of Nemp1 in intact NE. Note the lack of accumulation of either Lap2 (red trace) or chromatin (blue trace) in Nemp1 aggregates. Right panel: Maximum intensity projection (Max I) of the same cell showing the preferential accumulation of Nemp1aggregates near or at the edge of the NE opening.* ***E.*** *Maximum intensity projection of WT (top) or Nemp1 KO (bottom) erythroblasts undergoing NE opening. Note the absence of Nemp1 aggregates in Nemp1KO erythroblasts thereby confirming their specificity.*

***Video S1:*** *Non-tresholded 3D reconstruction from confocal planes of an erythroblast undergoing nuclear envelope opening*

***Video S2, S3****: 3D reconstruction from confocal planes (thresholded for Nemp1 intensity to emphasize Nemp1 aggregates, top panels) of erythroblasts undergoing a large (top) or small (bottom) nuclear envelope opening. Note the presence of Nemp1 aggregates near or close to NE openings.*
